# Hakai links m^6^A RNA methylation to immune regulation in colorectal cancer

**DOI:** 10.64898/2025.12.04.691578

**Authors:** Macarena Quiroga, Juan-José Escuder-Rodríguez, Andrea Iannucci, Victoria Suarez, Ivan Monteleone, Angélica Figueroa

## Abstract

**Background:** The m⁶A modification is the most abundant RNA epigenetic mark in colorectal cancer (CRC). In this study, we focused on Hakai, a methyltransferase writer–associated protein whose molecular function within the complex and link to CRC immune regulation remain poorly characterized.

**Methods:** We performed RNA-seq to assess transcriptomic changes after Hakai silencing in CRC models and MeRIP-seq to profile methylation alterations. Validation was carried out by Western blot, flow cytometry, ELISA, and RT-qPCR. Protein interactions within the writer complex were analysed by co-immunoprecipitation, and changes in protein localization by cell fractionation assays. Peripheral blood from healthy donors and lamina propria mononuclear cells from CRC patients were incubated with supernatants from Hakai-silenced CRC cells to evaluate effects on immune responses.

**Results:** Hakai silencing induced phenotypic changes in both monolayer and 3D CRC cultures, while its impact on the global transcriptome was limited. However, significant alterations in the methylation of immune response–related genes and reduced total m⁶A levels were observed. Hakai interacted with the writer-associated protein VIRMA, and its silencing altered the subcellular localization of METTL3. Moreover, conditioned media from Hakai-silenced CRC cells modulated the expression of immune response markers in patient-derived gut immune cells.

**Conclusions:** Hakai acts as a component of the m⁶A writer complex in CRC cells, influencing RNA methylation and the expression of immune-related genes. These findings suggest that Hakai silencing may enhance antitumour immune activity and represent a potential strategy to boost cancer immunity in colorectal cancer.

## BACKGROUND

In 2022, the estimated number of deaths due to cancer was estimated at 9.7 million, and of these, 900,000 deaths were due to colorectal cancer (CRC), ranking second as the highest mortality cancer, only behind lung cancer [1]. Beyond its common epithelial origin, CRC comprises a group of molecularly heterogeneous diseases characterized by diverse genomic and epigenomic alterations [2]. The epigenetic RNA methylation, especially the most abundant N6-methyladenine (m^6^A) modification [3,4], regulates mRNA stability, splicing, translation efficiency, and degradation, and has important implications for tumour biology and cancer therapy [5–7]. m^6^A is a reversible modification of mRNA, with a varied set of proteins that introduce (writers or m^6^A methyltransferases), recognize (readers), and remove (erasers or m^6^A demethylases) the methyl group. In cancer, m^6^A is critical for tumour initiation, progression, invasion, and metastasis. It also influences cancer stem cell pluripotency, differentiation, proliferation, migration, and tumour immunity. On the other hand, m^6^A can have tumour suppressor effects through the inhibition of specific oncogenes while promoting the expression of tumour suppressor genes [8]. The role of the m^6^A modification in cancer development is context dependent, and is influenced by the type of cancer, the expression and activity of m^6^A writers, erasers and readers, the cellular context and the tumour microenvironment [4].

m^6^A writers are S-adenosylmethionine-dependent methyltransferases (SAM) that perform N6-methylation by transferring a methyl group from the SAM cofactor to the adenosine substrate [9]. The m^6^A writer complex is a heterodimeric core comprised of enzymes METTL3 (Methyltransferase-like 3) and METTL14 (Methyltransferase-like 14), also called the m^6^A-METTL Complex (MAC), that was structuraly resolved in 2016 [10]; along with accessory proteins, organized in the m^6^A METTL-associated complex (MACOM), including WTAP, Hakai (CBLL1 gene), ZC3H13, VIRMA, and RBM15/15B, with a structure resolved with cryo-electron microscopy in 2022 [11].

The involvement of Hakai within the complex, is one of the less characterized from all the complex components so far. Hakai has been first and mostly described as the E3 ubiquitin-ligase responsible for ubiquitination of Src-phosphorylated E-cadherin, leading to its degradation that causes the disruption of cell-to-cell contacts, a process known as early as 2002 [12]. In this context, Hakai is known to form a homodimer required for its E3 ubiquitin-ligase activity, allowing the formation of an unusual phosphotyrosine-binding pocket involving its RING-finger domains, the Hakai-pY-binding (HYB) domain [13,14]. An early proteomics study proposed a nuclear complex involving Hakai’s interaction through its RING finger domain with WTAP in 2013 [15], but it was not until 2017 that Hakai was found as a component of the writer complex in plants [16]. This was followed in 2021 with a study in *Drosophila*, where it was shown to contribute to the stability of the writer complex [34].

Thus, Hakai’s involvement in the m^6^A methylation complex has opened an unexplored field regarding its potential role in RNA metabolism. As one of the less characterized members of the complex, research is needed for understanding the precise molecular mechanisms that underlie Hakai’s regulation of m^6^A methylation in CRC. This in turn would contribute to our understanding on how to modulate m^6^A in a context specific manner that may be beneficial for cancer patients.

## METHODS

### Cell culture

#### 1. Cell lines

Colorectal cancer epithelial cell lines HCT116 (CVCL_0291) and SW480 (CVCL_0546) from the American Type Culture Collection (ATCC) were cultured in DMEM (Thermo Fisher Scientific). HT29 (CVCL_0320) human colon adenocarcinoma cell line from Sigma-Aldrich was cultured in McCoy’s 5A medium (Thermo Fisher Scientific). The three cell lines were supplemented with 10% FBS (Corning) and 1% Penicillin/Streptomycin (5000U/mL; Thermo Fisher Scientific). Cell lines were periodically tested for *Mycoplasma* contamination and were used for a maximum of 3 months after thawing. Authentication of cell lines was performed at the Genomics platform of the Complutense University of Madrid (Spain). HT29 cell line with inducible silencing of Hakai was generated as previously reported [17]. Briefly, a lentiviral vector system for knocking-down Hakai expression (SMART vector Inducible Lentiviral shCBLL1) using doxycycline-inducible sh-RNA particles (Sigma-Aldrich) was acquired from Dharmacon, as well as an empty vector control. Cells were cultured in McCoy’s 5A medium with 1 μg/mL doxycycline for at least 72 hours to achieve optimal silencing, unless specified otherwise. For 3D tumourspheres cultures were stablished from the HT29 cell line with stable Hakai silencing (induced with doxycycline for 48 h). Cells were plated on ultra-low attachment 6-well plates (Corning) in DMEM/F-12 (Thermo Fisher Scientific) supplemented with factors that favour the formation of tumourspheres (**Supplementary Table 1**). Formation of tumourspheres was observed by contrasting phase microscopy using an Eclipse-Ti microscope (Nikon).

#### 2. Patients ans samples

Peripheral blood mononuclear cells (PBMCs) were obtained from 6 healthy subjects at the Tor Vergata University Hospital (Italy). Colon tissue samples were taken from macroscopically unaffected areas of **3** patients who underwent colon resection for sporadic CRC at the Tor Vergata University Hospital and used for the isolation of lamina propia mononuclear cells (LPMC). No patients received radiation therapy or chemotherapy prior to surgery. Human studies were approved by the local ethics committee. Additionally, human samples were collected from General and Digestive Surgery Service of the A Coruna University Hospital (Spain) for LPMC isolation. PBMC were also collected from samples from the Galician Blood, Organs and Tissues Agency (ADOS, Spain). Studies with human samples were approved by the local ethics committee. A total of 6 healthy subjects and 3 sporadic CRC patients samples were employed in this work. PBMCs extraction was carried out using density gradients with lymphoprep solution (Stemcell Technologies). LPMCs were obtained as indicated before [18]. In detail, mucosa fragments were incubated in HBSS with 1 mM EDTA (Thermo Fisher Scientific). Digestion was carried out with 200 μg/mL DNase I (Thermo Fisher Scientific) and 200 μg/mL Liberase^TM^ (Roche) for 1h at 37 °C. Cells were cultured in RPMI 1640 medium (Thermo Fisher Scientific), supplemented with 10% FBS, 1% Penicillin/Streptomycin, 2.5 μg/mL Amphotericin B (Thermo Fisher Scientific) and 10 μg/mL Gentamicin (Thermo Fisher Scientific).

### Plasmids, reagents, and antibodies

Plasmids pcDNA-Flag-Hakai and pcDNA 3.1 were kindly provided by Dr. Yasuyuki Fujita (Hokkaido University, Japan) and plasmid pTriEx 1.1-T2A-eGFP-Flag-Virilizer was kindly provided by Dr. Jianzhao Liu (Life Sciences Institute, Zhejiang University, China). The Hakai inhibitor, Hakin-1 [19], was used at 50 μM for 48 h. Antibodies used in this study are detailed in **Supplementary Table 2**.

### Western Blotting and immunoprecipitation

Protein extraction was performed with lysis buffer (20 mM Tris-HCl pH 7.5 [Sigma-Aldrich], 150 mM NaCl [Sigma-Aldrich], 1% Triton X-100 [Sigma-Aldrich], 10 μg/mL aprotinin [Sigma-Aldrich], 10 μg/mL leupeptine [Sigma-Aldrich] and 1 mM phenylmethylsulfonyl fluoride [Sigma-Aldrich]). For immunoprecipitation (IP), 10mM cysteine N-ethylmaleimide (Sigma-Aldrich) was added to the lysis buffer. NZYColour Protein I (NZYTech) marker was loaded to assess molecular weight. The chemiluminescent signal from the HRP-conjugated antibodies was detected with the Immobilon® Crescendo Western HRP (Millipore) substrate detection reagent using an Amersham Imager 600 imaging system (GE Healthcare).

### Quantitative Polymerase Chain Reaction (q-PCR) gene expression analysis

RNA extraction was performed with the TriPure Isolation (Roche) reagent. Complementary DNA (cDNA) synthesis was carried out through reverse transcription polymerase chain reaction (RT-PCR) using the NZY First Strand DNA synthesis kit for RT-PCR (NZY Tech). Template RNA was degraded using NZY RNase H (NZY Tech). The reverse transcription for Methylated RNA Immunoprecipitation Sequencing (MeRIP-Seq), was performed with the SuperScript™ VILO™ cDNA Synthesis Kit (Thermo Fisher Scientific). q-PCR was performed with the LightCycler® 480 SYBR Green I Master (Roche). The genes and primer sequences used in q-PCR are detailed in **Supplementary Table 3**. β-actin and GAPDH were used as house-keeping genes.

### Transcriptomic study by massive RNA sequencing

The transcriptomic study was carried out on HT29 cells with stable Hakai-silencing induced with doxycycline, obtained from monolayer or 3D-tumourspheres cultures. Extracted RNA was treated with DNase I (Thermo Fisher Scientific). RT-PCR was performed with the SuperScript™ VILO™ cDNA Synthesis Kit, in a library pre-loaded IonCode™ 96-well PCR plate. cDNA libraries were constructed using the Ion AmpliSeq™ Kit for Chef DL8 (Thermo Fisher Scientific) and the automated Ion Chef system, with the Ion AmpliSeq™ Transcriptome Human Gene Expression Panel (A31446). Libraries were loaded in the Ion 540™ Chip (Thermo Fisher Scientific). The sequencing was carried out in an Ion S5XL (Thermo Fisher Scientific), using the human genome version 19 (GRCh37) as reference. Three independent experimental replicates were performed for each condition.

### Bioinformatics analysis

RNAseq data was analysed using the Ion Torrent Suite software (Thermo Fisher Scientific). Differential expression analysis was performed in pairwise comparisons (each group corresponded to three independent experiments). A log-fold scale (LFC) shrinkage correction and dispersion estimation for low-count genes were applied. P-values were adjusted using the Benjamini–Hochberg method, and genes with adjusted p-value (FDR) < 0.1 and |LFC| > 2 were considered significant. Principal component analysis (PCA) was performed on normalized read counts transformed with the regularized logarithm (rlog) method. Functional annotation the differentially expressed genes was conducted using Gene Ontology (GO) (http://geneontology.org). The analysis of interactive networks was performed with STRING (http://string-db.org).

### Total m^6^A methylation of RNA quantification

Quantification of m^6^A methylation of RNA (from MeRIP-Seq) was performed using the m^6^A EpiQuik colorimetric kit (Epigentek), following the manufacturer’s protocol. A negative control (m^6^A-free RNA) and a positive control (m^6^A-modified oligonucleotides normalized to 100% signal) were used to normalize sample data. Absorbance was read on a plate reader at 450 nm.

### Methylated RNA Immunoprecipitation Sequencing (MeRIP-seq)

Hakai-silenced HT29 cells were used for epitranscriptomic analysis. RNA fragmentation was performed with fragmentation buffer (100mM Tris-HCl [pH 7.0] [Sigma-Aldrich], 10mM ZnCl_2_ [Sigma-Aldrich], at 94 °C for 5 min. Correct RNA fragment size was verified by agarose-gel electrophoresis with Sybr Safe DNA Stain (Thermo Fisher Scientific) with a control 100 bp DNA Ladder (Sigma-Aldrich). For IP, RNA fragments were pooled and incubated in RNase-free water containing 300 U RNasin (Thermo Fisher Scientific), 2mM vanadyl-ribonucleoside complex (Sigma-Aldrich), 12.5 µg antibody against m^6^A (Synaptic Systems), 5X IP buffer (50 mM Tris-HCl [Sigma-Aldrich], 750 mM NaCl [Sigma-Aldrich], and 0.5% [vol/vol] Igepal CA-630 [Sigma-Aldrich]), for 2 hours at 4 °C with constant mixing. Immunoglobulin G (IgG) was used as negative control in the IP. Protein *A* agarose beads (Santa Cruz) were incubated with the sample containing the m^6^A antibody and with control IgG. Elution was achieved with elution buffer (1X IP buffer, 6.7 mM m^6^A [Selleckchem] and 280 U RNasin, in RNase-free water), for 1 hour at 4 °C. cDNA synthesis was performed by RT-PCR. IgG libraries were sequenced for normalization correction.

### Transfections

Lipofectamine^TM^ 2000 (Invitrogen) and the appropriate plasmid were diluted separately in Opti-MEM^TM^ (Thermo Fisher Scientific) and mixed before replacing cell culture media. Cells were collected 48 h after transfection. Hakai was silenced with small interfering ribonucleic acid Hakai-1 oligonucleotide sequence (5’ CTCGATCGGTCAGTCAGGAAA), following the same protocol as with plasmids. Mission Universal Non-coding siRNA (Sigma-Aldrich) was used as negative control.

### Inhibition Assays

Protein half-life experiments were performed using the protein synthesis inhibitor cycloheximide (CHX) (Sigma-Aldrich). HCT116 cells were transfected with pcDNA-Flag-Hakai. 48 h after transfection, 10 μg/mL CHX was added and incubated for the specified times. Actinomycin D (ActD, Sigma-Aldrich) was used to assess RNA stability. HT29 cells were induced for Hakai-silencing with doxycycline for 72 h. Then, 10 μg/mL ActD was then added and incubated for the specified times. The global methylation inhibitor 3-deazaadenosine (3-DZA) (Sigma-Aldrich) was used to inhibit RNA methylation. HT29 cells were induced for Hakai-silencing with doxycycline for 48 h. Then, 10 μg/mL 3-DZA was added with an incubation of 48 h.

### Flow cytometry

For flow cytometry, PBMCs and LPMCs were cultured for 24 hours in supplemented RPMI 1640 medium. The medium was discarded and the supernatant from either HT29 cells with Hakai-silencing or HT29 control was added. After 24 hours, the cells were collected. Cell suspensions were incubated with Fc receptor blocking solution (Human TruStain FcX™, BioLegend) for 10 minutes at room temperature, before staining. To evaluate cell viability, Live/Dead staining (Invitrogen) was added and incubated on ice for 10 minutes in darkness. Specific antibodies were used for each protein target (**Supplementary Table 2**). For extracellular proteins, cells were incubated 30 minutes at 4 °C in darkness with the specific antibody. For intracellular proteins, fixation with eBioscience™ fixation buffer (Thermo Fisher Scientific) and permeabilization with eBioscience™ permeabilization buffer (Thermo Fisher Scientific) were performed. Detection was carried out on a CytoFLEX cytometer (Beckman Coulter).

### Enzyme-linked immunosorbent assay (ELISA)

For ELISA assays, Human IL-8/CXCL8 (DuoSet® R&D BioSciences) and Human CXCL1/GRO Alpha (DuoSet® R&D BioSciences) kits were used. Absorbance was measured using the NanoQuant Infinite® M200 PRO spectrophotometer (Tecan, Switzerland) at 450 nm with a reference absorbance for correction at 540 nm. Determinations were made in duplicate.

### Immunoprecipitation (IP) assay and cell fractionation assay

HCT116 cells were transfected 48 h after establishment of cell culture, using plasmids pcDNA-FLAG-Hakai and pcDNA3.1 as negative control. *Protein A* agarose beads (Santa Cruz) were incubated with Hakai’s antibody (Invitrogen). As a negative control, Hakai antibody was replaced with IgG. Fractionation was performed using the Subcellular Protein Fractionation Kit for Cultured Cells (Thermo Fisher Scientific) on HT29 cells with Hakai-silencing and control conditions.

### Statistical analysis

The statistical significance of the data was determined using Student’s t-test analysis when comparing two groups and by ANOVA analysis for more than two groups. The significance between the experimental groups indicated in the figures is shown as *p < 0.05, **p < 0.01 and ***p < 0.001. The results obtained are expressed as mean ± SEM. Graphical representations, as well as statistical analyses, were performed using GraphPad Prism 9.0.2 (GraphPad Software).

## RESULTS

### Hakai exerts a limited impact on the transcriptional landscape of CRC monolayer and 3D tumoursphere cultures

Although the post-translational regulatory mechanism of Hakai is well established, little is known about its potential role at the transcriptional or post-transcriptional level in CRC. To identify Hakai-modulated transcriptional pathways we conducted RNA sequencing (RNA-seq) in CRC cells cultured under both monolayer and tumourspheres conditions, in the presence or absence of Hakai. With this approach we sought to obtain a more comprehensive view of Hakai’s role in the tumour biology of CRC. Hakai silencing was confirmed by Western blot analysis in monolayer cultures of HT29 cells (**Figure 1A**). Control cells showed an epithelial phenotype, evidenced by its growth as a monolayer and the formation of tightly connected cell clusters, however, Hakai silencing clone showed a prominent three-dimensional cell sheet organization, presenting cells more closely adhered to each other compared to control cells (**Figure 1B**). Gene expression profiling by RNA-seq, followed by PCA, revealed a clear separation between the experimental groups along the first two principal components (PC1 and PC2) (**Figure 1C**). A total of 115 genes were found to be differentially expressed in Hakai silencing HT29 cells compared to control HT29 cells. However, only the silenced gene, CBLL1, exceeded the established significance threshold (**Figure 1D-F**), suggesting that the overall transcriptomic impact of Hakai silencing is limited and may not strongly affect global gene expression under these conditions. Nevertheless, given the restrictive significance threshold adopted, we performed a STRING analysis with the 115 differentially expressed genes (DEGs). This analysis revealed several protein-protein interaction (PPI) networks of the proteins encoded by differentially expressed genes due to Hakai silencing (**Figure 1G**). These included a cancer-related protein network (GRB2, PIK3R2, CCND1, SMAD3, CASP3, RPS6KB2, FGF19, ITGAV, EGLN3, IL6ST, NFKB2 and IFNGR1) and a colorectal cancer-related protein network (GRB2, PIK3R2, CCND1, SMAD3, CASP3 and RPS6KB2); a protein network involved in viral carcinogenesis (GRB2, PIK3R2, CCND1, IRF7, CASP3, H4C6, H2BC21, H2BC15, H2BC5, IL6ST, and NFKB2), and a protein network involved in viral response (DTX3L, IRF1, IRF7, DDX58, IFIH1, IRF5, SAMHD1, IFIT3, OAS3, IFIT2, IFIT5, DHX58, SMAD3, IFNGR1, and EXT1). Moreover, Gene Ontology enrichment analysis of biological processes (**Figure 1H**), also highlighted enrichment of immune response–related categories. Remarkably, enrichment of signalling pathways associated with antiviral response and innate immunity, such as the MDA-5 signalling pathway and the upregulation of interferon alpha and beta production, was observed. Regarding cell localization (**Figure 1H**), Hakai regulated genes showed significant enrichment in components associated with extracellular trafficking,. When evaluating molecular functions (**Figure 1H**), an enrichment of RNA binding-related functions was found. We then analysed the effects of Hakai silencing on the 3D tumourspheres model (**Figure 1I**). As shown in **Figure 1J**, Hakai silencing significantly reduced tumoursphere size. However, variability between the two experimental conditions was less evident than in the monolayer conditions, using the first two principal components of the analysis (**Figure 1K**). RNA-seq analysis identified nine DEGs in tumourspheres derived from Hakai silencing HT29 cells compared to control HT29 tumourspheres (**Figure 1L-N**). Among these, two showed significant differences: CCL5 (encoding a chemokine), and CBLL1 (Hakai, silence-targeted gene) (**Figure 1N**). These findings reinforce the notion that Hakai silencing exerts a moderate impact on the global gene expression profile of CRC cells. Nevertheless, STRING analysis was performed to explore the predicted interaction network of proteins encoded by the identified DEGs, revealing enrichment in processes related to the downregulation of viral genome replication (**Figure 1O**). Taken together our results idicate that Hakai silencing has a modest effect on the transcriptome in both monolayer cultures and 3D tumourspheres of CRC cells.

**Figure 1.**
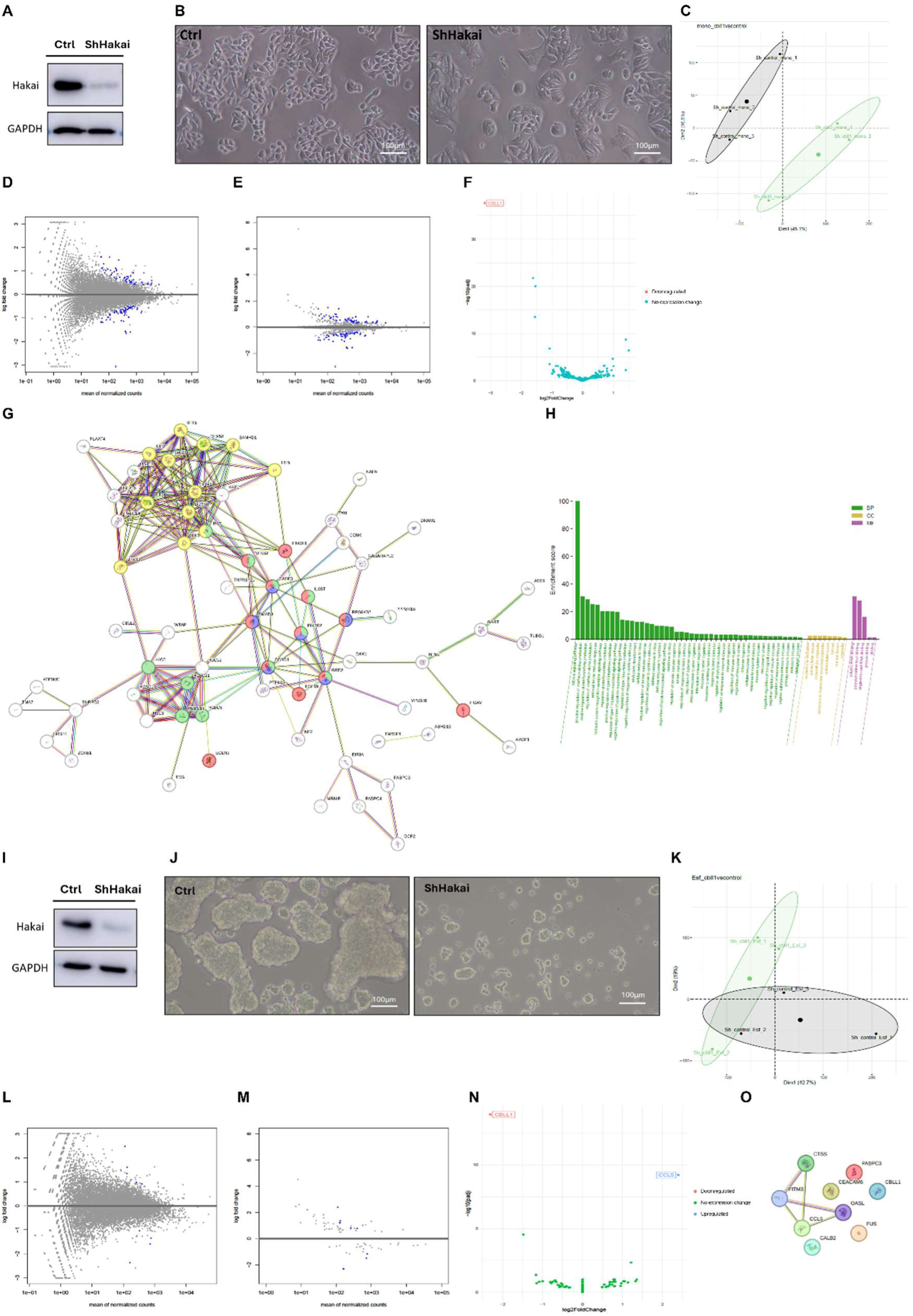
Hakai silencing induces phenotypical changes in colorectal cancer cells but exerts a limited impact on their overall transcriptional landscape. (A) Western blot showing Hakai downregulation with a Hakai-directed sh-RNA in monolayer cultures of HT29 cells. (B) Phase contrast microscopy images of HT29 cells of the control conditions (left) and under Hakai-silencing conditions (right). (C) Principal component analysis (PCA) of gene expression profiling by RNAseq. (D) Mplot of the log fold change (LFC) estimates for all genes represented in the libraries based on normalized count values. (E) Mplot of the LFC estimates after false discovery rate (FDR) correction. (F) Volcano plot of differentially expressed genes (DEGs) between Hakai silencing and control conditions with an FDR < 0.1 and a LFC > 2. (G) STRING Protein-Protein Interaction (PPI) network of the proteins encoded by the DEGs. Coloured nodes represent cancer-related networks (red), colorectal cancer-related networks (blue), viral carcinogenesis networks (green) and viral response networks (yellow). Protein nodes with no connection are hidden. (H) Gene Ontology (GO) enrichment analysis of biological processes (BP), cellular compartment (CC), and molecular function (MF). (I) Western blot showing Hakai downregulation with a Hakai-directed sh-RNA in 3D tumourspheres. (J) Phase contrast microscopy images of 3D HT29 tumouspheres in control (left) and Hakai silencing conditions (right). (K) PCA of the gene expression profiling by RNA-seq in 3D HT29 tumourspheres. (L) Mplot of the LFC estimates for all genes represented in the libraries, based on normalized count values. (M) Mplot of LFC estimates after FDR correction. (N) Volcano plot of DEGs between Hakai silencing and control conditions in 3D tumourspheres with an FDR < 0.1 and a LFC > 2. (O) STRING PPI network for the nine DEGs indentified in 3D tumouspheres with Hakai silencing.

### Hakai regulated m^6^A methylation of RNA in colorectal cancer cultures

Our results indicate that Hakai silencing has limited impact on the overall transcriptome of CRC, as revealed by the RNA-seq data from monolayer and 3D tumourspheres models. However, previous studies have demonstrated the participation of Hakai in the regulation of m^6^A methylation in other species and cell lines. To assess whether Hakai influences global RNA methylation in CRC cells, total m^6^A levels were measured in HT29 cells in presence and absence of Hakai (**Figure S1A**). Our results showed a 25% reduction in total m^6^A RNA levels upon Hakai silencing (**Figure 2A**). We next aimed to analyse the expression profile of methylated genes using MeRIP-seq in HT29 cells with or without Hakai, providing insight into how Hakai may influence the post-transcriptional regulation of genes (**Figure 2B**). Upon Hakai silencing (**Figure S1B**), extracted RNA was fragmented into approximately 100 nucleotides-sized fragments (**Figure S1C).** PCA calculation revealed no evident separation between the two experimental conditions along the first two principal components (PC1 and PC2) (**Figure 2C**). Thus, the second and third principal components (PC2 and PC3) were also plotted to better visualize variation between groups (**Figure 2D**). A total of 56 genes were found differentially methylated in Hakai silencing HT29 cells compared to control cells. When silencing Hakai, 31 genes (55%) showed a down-regulation in their methylation, while 25 (45%) were up-regulated. To investigate interactions among differentially methylated genes, a STRING PPI network analysis was performed (**Figure 2E**). Interestingly, this analysis revealed PPI networks related to inflammatory and immune responses, including the interaction between cytokines and cytokine receptors (TNF, CXCL8, CXCL1, CSF1 and IFNβ1) (**Figure 2F**), the TNF signalling pathway (TNF, PIK3CB, TNFAIP3, CXCL1, CSF1 and IFNβ1) (**Figure 2G**) and IL-17 (TNF, TNFAIP3, CXCL1 and CXCL8) (**Figure 2H**).

**Figure 2.**
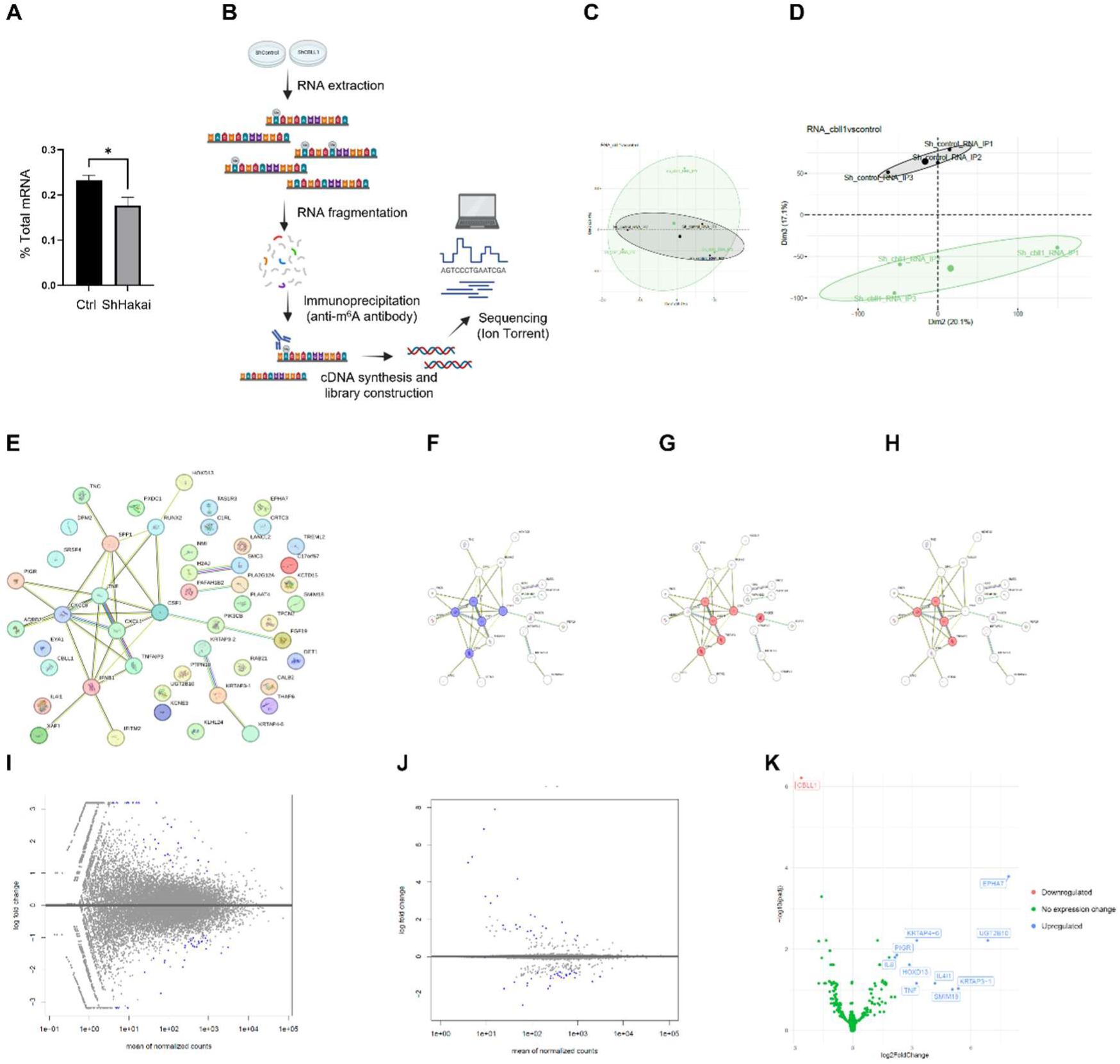
Effect of Hakai on m^6^A RNA methylation in HT29 cells. (A) Total m^6^A RNA levels in control conditions and upon Hakai silencing in HT29 cells. Data are expressed as normalized mean ± SEM of three independent experiments. (B) Methylation RNA immunoprecipitation sequencing (MeRIP-seq) workflow. (C-D) Principal component analysis (PCA) on principal components 1 and 2 (C) and 2 and 3 (D). Sample data from three independent experiments. (E) STRING PPI network of the 56 proteins encoded by genes differentially methylated upon Hakai silencing. (F) PPI network highlighting cytokines and cytokine receptors (in blue). Nodes with no connection are hidden. (G) PPI network highlighting TNF signalling pathway (in red). (H) PPI network highlighting IL-17 signalling pathway (in red). (I) Mplot of the LFC estimates for all genes represented in the libraries based on their normalized count values and (J) Mplot estimates of the LFC after FDR correction. (K) Volcano plot showing genes with different methylation between control and Hakai-silencing conditions.

To identify genes with significant methylation changes between experimental conditions, differential expression analysis was performed.. Eleven genes (20%) had statistically significant differences: 10 were upregulated, while only the silenced target gene, CBLL1, showed decreased methylation (**Figure 2I-K**).

### Hakai regulates m^6^A methylation of specific genes

Of the 55 differentially methylated genes (excluding Hakai) identified in Hakai-silenced HT29 cells compared to control cells, 15 were selected for validation by RT-qPCR. Selection criteria inclued the magnitude of the change in their expression (LFC) and the statistical significance (FDR). Given the limited number of genes meeting the significance threshold, additional genes with relevant biological functions were also included, despite not surpassing the restrictive predefined statistical cutoff. Validation of the selected genes by RT-qPCR upon Hakai silencing (**Figure S2A**) in HT29 cells showed an increase in the expression levels of methylated CXCL1, IFNβ1, PIGR, TNF, CXCL8, EPHA7 and HOXD13 compared to control cells (**Figure 3A**), in agreement with the results obtained from the identification by MeRIP-seq. Moreover, methylated PTPN18 and FGF19 levels decreased upon Hakai silencing in HT29 (**Figure 3B**), also consistent with the results obtained in the MeRIP-seq analysis. However, no significant changes were validated fog PAFAH1B2, RAB21, IL4I1, DET1, PIK3CB and TREML2 genes (**Figure 3C**). These results show that many of the genes selected for RT-qPCR validation displayed significant methylation changes upon Hakai silencing, consistent with the MeRIP-seq findings.

**Figure 3.**
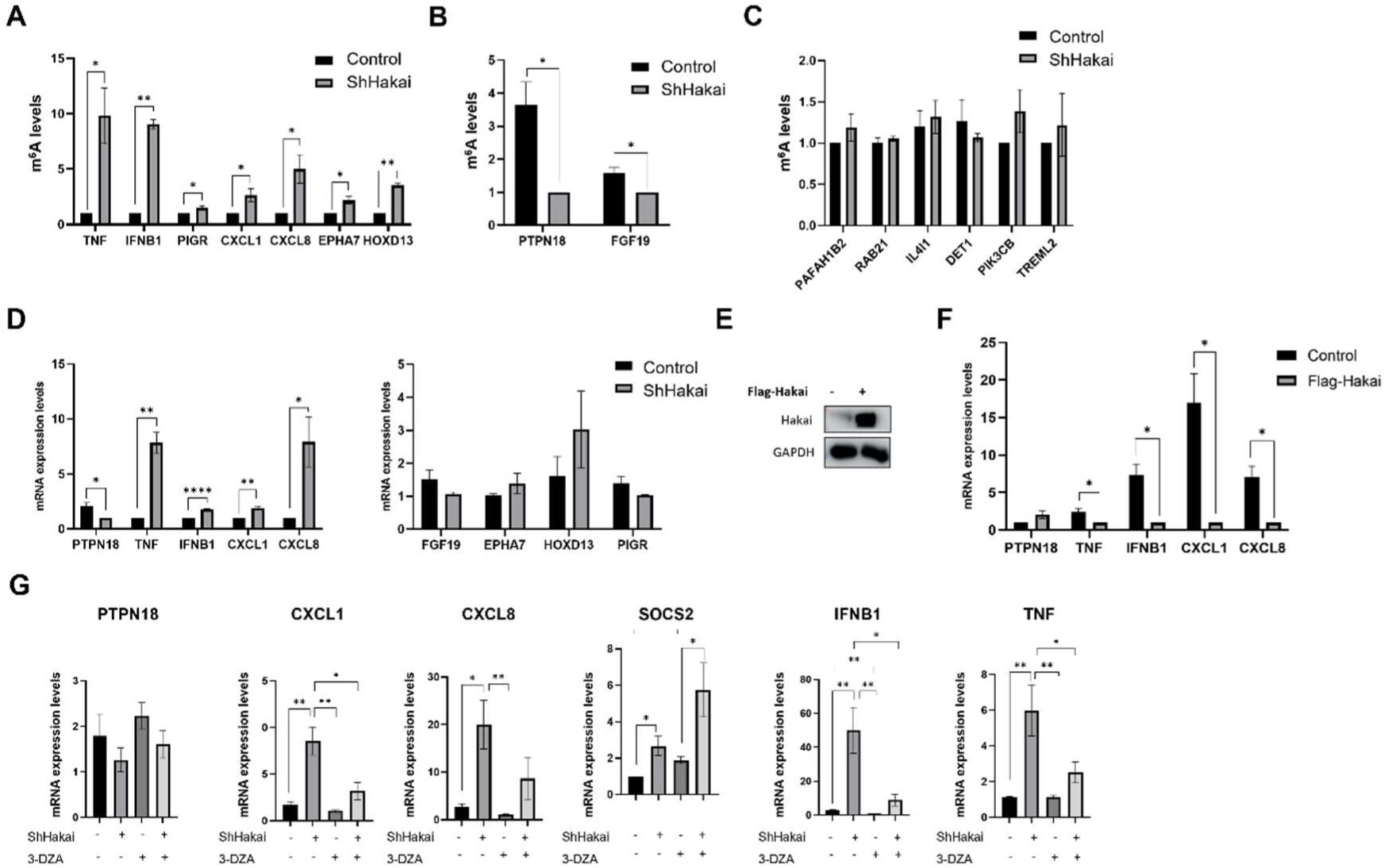
Hakai involvement in the m^6^A methylation of specific genes. RT-qPCR analysis of upregulated (A), downregulated (B) and non-modulated (C) m^6^A-immunoprecipitated genes, under control and Hakai-silencing conditions. (D) RT-qPCR analysis of differential (left) and non-regulated (right) gene expression of the selected genes. (E) Western blot showing Hakai overexpression following transfection with pcDNA-FLAG-Hakai plasmid. (F) RT-qPCR analysis of selected gene under control and Hakai-overexpression conditions. (G) RT-qPCR analysis of selected genes after m^6^A methylation blockade with 3-DZA assay in control and Hakai-silencing conditions. m^6^A levels and mRNA expression levels are expressed as normalized mean ± SEM of three independent experiments. * p < 0.05; ** p < 0.01; *** p < 0.001; **** p < 0.0001.

To investigate the potential involvement of the m^6^A methylation modification in the selected genes modulated by Hakai, we first analysed the total mRNA expression levels of these genes in presence or absence of Hakai (**Figure S2B**). As shown in **Figure 3D**, Hakai silencing significantly reduced PTPN18 mRNA expression, while increasing TNF, IFNB1, CXCL1, and CXCL8. In contrast, no significant changes were observed in the expression of FGF19, EPHA7, HOXD13, and PIGR.

We next investigated PTPN18, TNF, IFNB1, CXCL1, and CXCL8 gene expression following Hakai overexpression. To this end, SW480 cells were transfected with the plasmid pcDNA-Flag-Hakai (**Figure 3E**) and target gene expression was evaluated by RT-qPCR. Hakai overexpression lead to reduced mRNA levels of TNF, IFNB1, CXCL1, and CXCL8 in SW480 cells, whereas no significant changes in PTPN18 gene expression were detected (**Figure 3F**). These results are consistent with those obtained under Hakai silencing, supporting the notion that Hakai negatively regulates inflammatory gene expression, although its effect on PTPN18 appears limited in this model.

Next, to assess whether Hakai modulates gene expression through m^6^A-dependent mechanisms, we evaluated the effect of Hakai silencing combined with the global methylation inhibitor 3-DZA (**Figure S2C**) by RT-qPCR, with SOCS2 as a positive control. As shown in **Figure 3G**, the upregulation of TNF, IFNB1, CXCL1, and CXCL8 mRNA observed upon Hakai silencing was reversed by 3-DZA treatment, supporting the role of Hakai in regulating these genes through m^6^A methylation. In contrast, no significant changes were observed in PTPN18 expression. As expected, SOCS2 mRNA levels increased following 3-DZA treatment, and this effect was further enhanced by Hakai silencing.

### Hakai-mediated m^6^A methylation of immune-related genes increases protein levels without affecting transcript stability

Based on our previous findings, which showed changes in the expression of TNF, IFNB1, CXCL1 and CXCL8 upon Hakai silencing and overexpression, we next investigated whether m^6^A methylation could influence the RNA stability of these transcripts. To this end, we performed RNA stability assay using ActD, a transcriptional inhibitor that blocks *de novo* RNA synthesis. As shown in **Figure 4A**, no significant differences in mRNA half-life were observed between control and Hakai-silencing conditions (**Figure S3A**) indicating that Hakai-mediated m^6^A-dependent mechanisms do not affect the stability of these transcripts.

**Figure 4.**
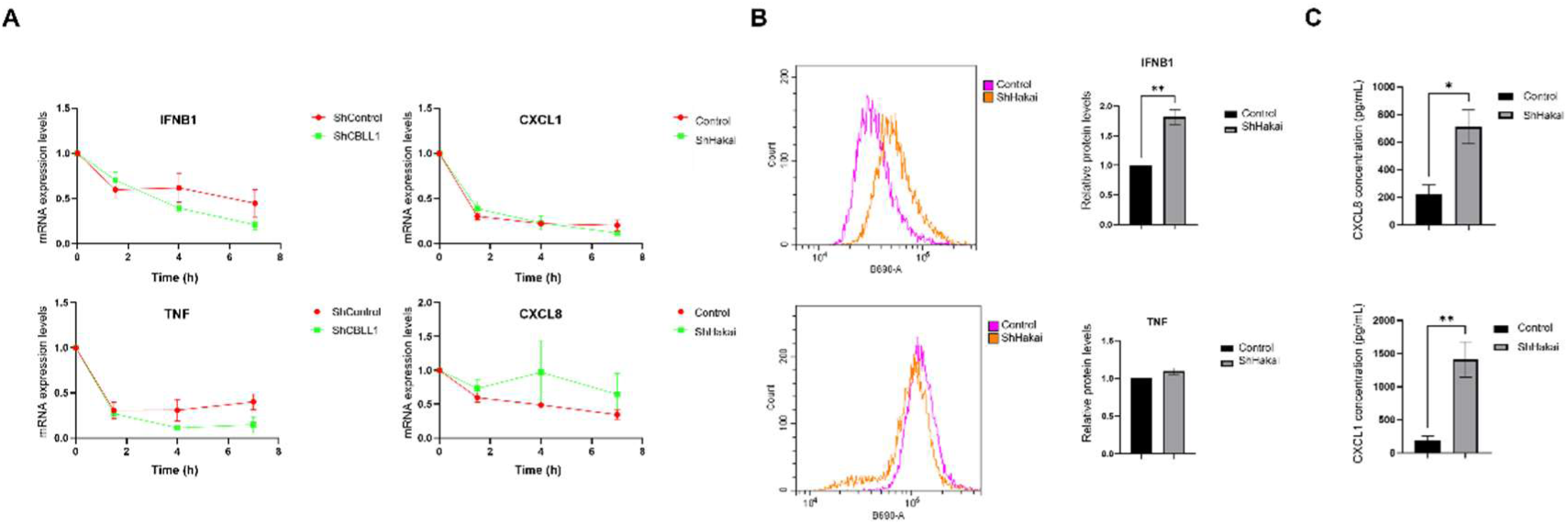
Effects of Hakai on transcript stability and protein levels. (A) RT-qPCR analysis of mRNA levels of Hakai-regulated genes under control and Hakai-silencing conditions. Data are expressed as normalized mean ± SEM of three independent experiments. (B) Flow cytometry analysis of IFNβ1 and TNFα protein expression. Relative protein expression levels are expressed as normalized mean ± SEM of three independent experiments. (C) ELISA measurement pf CXCL8 and CXCL1 protein levels. Protein concentrations are expressed as mean ± SEM of three independent experiments. * p < 0.05; ** p < 0.01.

Next, we investigated the impact of Hakai on protein expression levels. Hakai silencing (**Figure S3B**) increased IFNβ1 protein levels, whereas TNFα expression remained unchanged, as revealed by flow cytometry using specific antibodies (**Figure 4B**). Consistently, Hakai silencing (**Figure S3C**) also increased CXCL8 and CXCL1 protein levels as shown by ELISA (**Figure 4C**).

### Hakai incluence on VIRMA expression and METTL3 localization

To investigate potential interactions between Hakai and the m^6^A writer complex MAC and MACOM proteins, a co-immunoprecipitation assay was performed. At endogenous levels, no interaction was detected between Hakai and METTL3, METTL14, or WTAP (data not shown), whereas an interaction with VIRMA was observed (**Figure 5A**). Under overexpression conditions, consistent with these findings, Hakai again interacted with VIRMA but not with METTL3, METTL14, or WTAP (**Figure 5B**).

**Figure 5.**
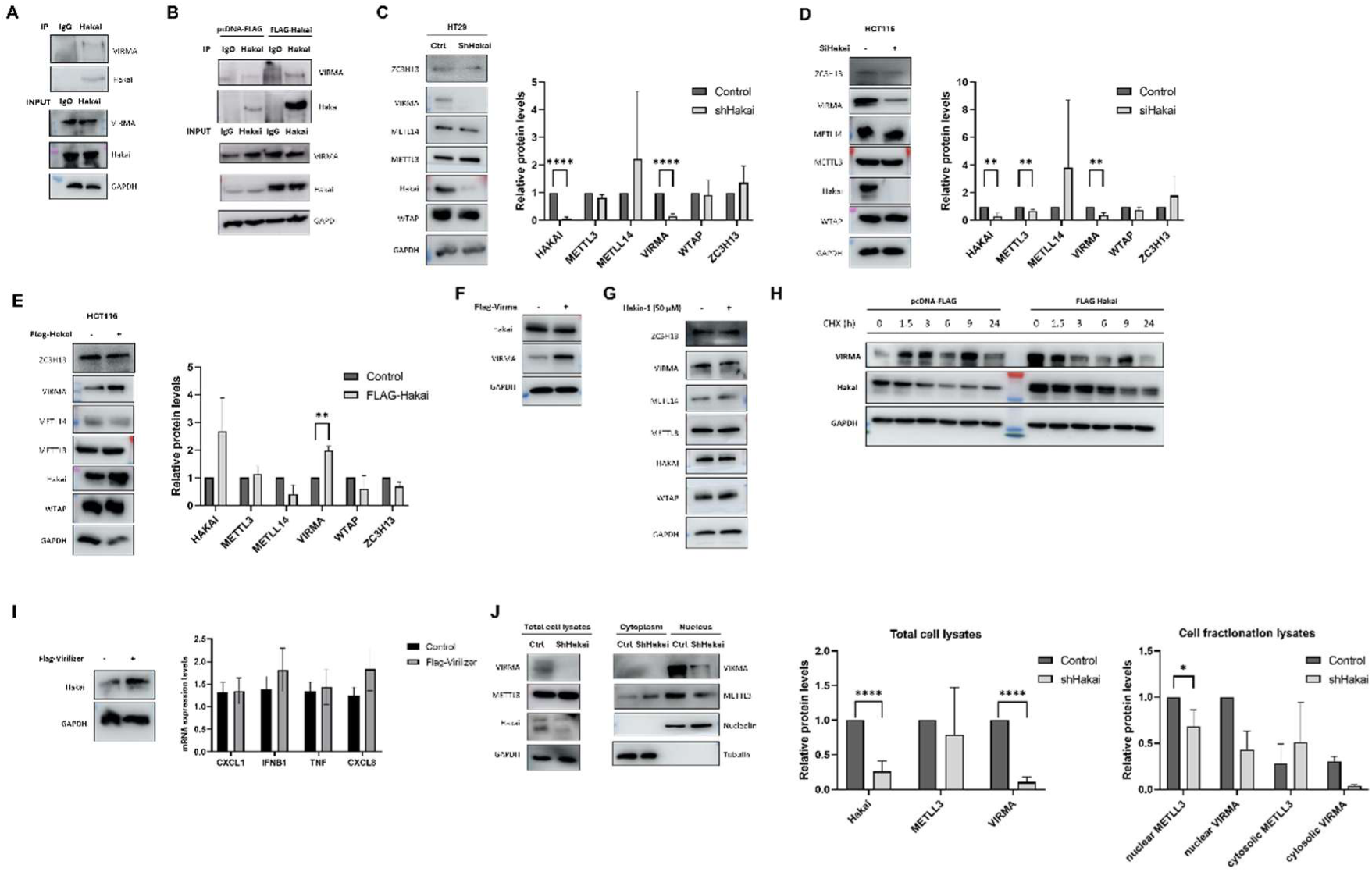
Interaction of Hakai with the m^6^A writer-associated complex. (A) Co-immunoprecipitation of endogenous Hakai with endogenous VIRMA in HCT116 colorectal cancer cells. (B) Co-immunoprecipitation of Hakai with endogenous VIRMA in Hakai-overexpressing HCT116 cells. (C-D) Western blot analysis of the effect of Hakai silencing on MAC and MACOM proteins in HT29 (C) and HCT116 cells (D). Data are expressed as mean relative protein expression ± SEM from three independent experiments. (E) Western blot analysis of the effect of Hakai overexpression on MAC and MACOM proteins in HT29. (F) Western blot analysis of the effect of VIRMA overexpression on Hakai protein levels in the HCT116 cells. (G) Western blot analysis of the effect of Hakin-1–mediated inhibition of Hakai E3 ubiquitin ligase activity on MAC and MACOM protein expression. (H) Western blot analysis of VIRMA protein stability in HCT116 cells following CHX treatment under Hakai-overexpression conditions. (I) RT-qPCR analysis of mRNA expression levels of Hakai-modulated trasncripts in VIRMA overexpression conditions. mRNA expression levels are expressed as normalized mean ± SEM of three independent experiments. (J) Western blot analysis of the subcellular location of METLL3 and VIRMA under Hakai-silencing conditions. Data are expressed as normalized mean ± SEM of three independent experiments. * p < 0.05; ** p < 0.01; *** p < 0.001; **** p < 0.0001.

Next, we investigated the molecular role of Hakai within the m^6^A methyltransferase complex. Given that Hakai regulates of protein expression, we evaluated the effect of Hakai silencing on of several complex components. Hakai silencing significantly reduced VIRMA protein levels in both HT29 (**Figure 5C**) and HCT116 (**Figure 5D**) cells, whereas WTAP, ZC3H13, METTL3, and METTL14 were unaffected. Conversely, Hakai overexpression (**Figure 5E**) increased VIRMA protein levels, without altering the expression levels of other writer complex components. Next, to determine whether the regulatory relationship between Hakai and VIRMA is reciprocal, we studied the impact of VIRMA overexpression on Hakai protein levels. VIRMA overexpression did not alter Hakai protein levels (**Figure 5F**), indicating that regulation is unidirectional: Hakai modulates VIRMA expression, whereas VIRMA does not influence Hakai. Since Hakai is a well-characterized E3 ubiquitin-ligase in mammals, we hypothesized that its ubiquitination activity might influence the stability of the writer complex components. We explored this hypothesis using Hakin-1, a specific inhibitor of Hakai designed to bind the HYB domain responsible for its E3 ligase activity. [18]As shown in **Figure 5G**, Hakin-1 treatment did not alter the levels of any of the tested proteins, including VIRMA. These results suggest that Hakai does not regulate the m^6^A writer complex through its E3 ubiquitin-ligase activity. Next, to test whether Hakai directly regulated VIRMA stability at the post-translational level, we measured VIRMA protein half-life under conditions of Hakai overexpression and protein synthesis inhibition with cycloheximide CHX. As shown in **Figure 5H**, Hakai overexpression did not alter the half-life of VIRMA in HCT116 cells. These findings further support that, under the experimental conditions employed, Hakai does not directly affect the post-translational stability of VIRMA.

Following these results, we investigated whether Hakai regulates target genes directly or indirectly via VIRMA. To this end, we evaluate the expression of TNF, IFNB1, CXCL1 and CXCL8 genes by RT-qPCR under VIRMA overexpression conditions. As shown in **Figure 5I**, VIRMA overexpression did not alter the expression of the genes studied in SW480 cells. These dfindings indicate that the modulation of TNF, IFNB1, CXCL1, and CXCL8 observed upon Hakai silencing and/or overexpression is not mediated by VIRMA levels and instead reflects a VIRMA-independent regulatory role of Hakai on these target genes.

Although Hakai silencing or overexpression did not affect the expression levels of METTL3, the catalytic core of the m^6^A writer complex, we observed a reduction in total m^6^A levels upon Hakai silencing. Given that m^6^A methylation primarily occurs in the nucleus, we hypothesized that Hakai may regulate this process by influencing the subcellular localization of key writer complex components. Cellular fractionation revealed that VIRMA is restricted to the nucleus and its expression decreases upon Hakai silencing, consistent with our previous findings. Under control conditions, METTL3 is predominantly nuclear, however, in the absence of Hakai, a notable fraction of the protein is redistributed to the cytoplasm (**Figure 5J**), suggesting that Hakai influences METTL3 nuclear–cytoplasmic distribution.

### Hakai regulates proinflammatory and cytotoxic gene expression in gut immune cells

To assess the effect of Hakai on the tumour microenvironment, PBMCs were incubated with supernatants from Hakai-silenced HT29 culture (**Figure S4**), and the CD45^+^ population (hematopoietic nucleated cells) was analysed by flow cytometry. Within this population, granzyme (natural killer [NK] and T cytotoxic cells marker) protein levels was analysed. Hakai silencing lead to a significant increase of granzyme expression in the CD45^+^ cells (**Figure 6A**), suggesting an increase, activation of cytotoxic immune cells. To further characterize the changes to immune response in the proximity of the tumours, lamina propria mononuclear cells, LPMCs were incubated with supernatants from Hakai-silenced HT29 culture, and the gene expression of pro-inflammatory, anti-inflammatory molecules were analysed by RT-qPCR. As shown in **Figure 6B**, Hakai silencing significantly increased IFNG, PRF1, and GZMB expression in LPMCs, in addition, although not statistically significant, there seems to be a mild increase in TNF and CXCL1, both of which can contribute to anti-tumour immune activation, together with a decrease in the immunosuppressive cytokines IL10 and TGFβ. Overall, these results suggest that Hakai silencing promotes a shift in the tumour microenvironment towards enhanced cytotoxicity and a pro-inflammatory, anti-tumour immune response.

**Figure 6.**
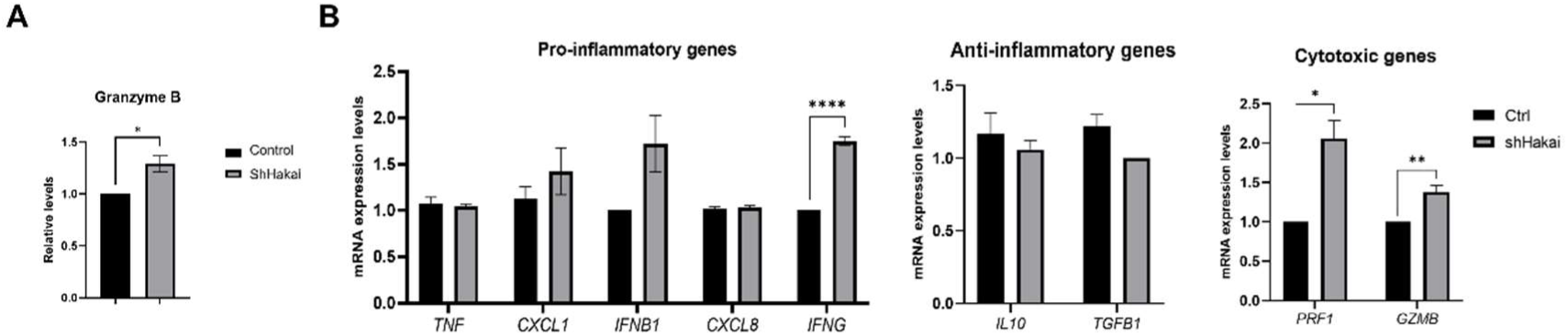
Effects of Hakai silencing on immune cell activation. (A) Flow cytometry analysis of of granzyme protein expression in the CD45^+^ PBMCs incubated with supernatants from control or Hakai-silencing conditions. Data are expressed as normalized mean ± SEM of three independent experiments. (B) RT-qPCR gene expression in LPMCs incubated with supernatants from control or Hakai-silenced HT29 cells. mRNA expression levels are expressed as normalized mean ± SEM of three independent experiments. * p < 0.05; ** p < 0.01; *** p < 0.001; **** p < 0.0001.

## DISCUSSION

It is well established that Hakai plays a key role in the EMT process by mediating the ubiquitination and degradation of src-phosphorylated E-cadherin at intercellular contacts, facilitating the loss of cell-cell adhesion and promoting cell migration and invasion [20,21]. Hakai’s relevance in EMT and in tumour metastasis has been the driving force behind most of the research focused on this protein [22]. However, in recent years, the potential role of Hakai in the nucleus has emerged. Recently, DEAD box helicase 39B (DDX39B) has been found to be responsible for the recruitment of both Src and Hakai to E-cadherin in cell-cell contacts [23]. Nonetheless, DDX39B had been mostly described as a nuclear mRNA-regulating protein, and this cytoplasmic role involving protein-protein interactions has been largely unknown. Similarly, Hakai is found in both the nucleus and the cytoplasm, showing cellular compartment-specific functions [24]. In contrast to DDX39B, Hakai’s function is mostly described in the context of posttransalational regulation, whereas its possible role in transcriptional regulation remains unexplored [25]. To contribute in filling this knowledge gap, we performed a comparative RNA-seq in CRC cells with and without Hakai (CBLL1 gene) expression. We employed the HT29 colorectal cancer cell line and a 3D tumorsphere model derived from it, which promotes cancer stem cell (CSC) survival and enrichment [26]. Other E3 ubiquitin-ligases such as FBXW7, NEDD4 or Skp2, have been shown to regulate CSCs traits, EMT and tumour self-renewal [27–29]. Our group recently demonstrated that Hakai silencing in CRC tumourspheres reduced their number and size and downregulated CSC markers (LGR5, Nanog, Klf4), supporting its role in CSC acquisition [30]. However, RNA-seq analysis revealed only modest transcriptome changes upon Hakai silencing (Figure 1), suggesting that Hakai is not a major regulator of global gene expression in CRC monolayers or 3D cultures. However, enrichment analysis revealed networks associated to carcinogenesis, immunity, vesicle trafficking and RNA regulation.

Multiple studies have linked Hakai to the m^6^A methylation process and the writer complex. In *Arabidopsis*, *Drosophila*, and human cancer cell lines (such as HeLa cells), Hakai silencing decreases total m^6^A levels by 35%, 50%, and 23%, respectively [16,31–33]. Our results in the HT29 CRC cell line are consistent with these findings (∼25% reduction, (Figure 2), supporting a critical role for Hakai within the evolutionarily conserved m^6^A complex as a positive regulator of m^6^A RNA methylation. However, the precise molecular mechanisms by which Hakai contributes to methyltransferase activity are unknown. Given that METTL3 is the only component of the writer complex with confirmed catalytic activity [34], it is plausible that Hakai contributes stabilizing of complex components or modulating its substrate specificity and/or RNA-binding affinity. Co-immunoprecipitation in the CRC cell line revealed an interaction between Hakai and VIRMA, but no interaction with other components (Figure 5). Thus, Hakai’s association with the m^6^A writer complex in CRC is likely mediated through the MACOM complex, specifically via VIRMA, rather than through the catalytic core. In pararell, we observed that Hakai silencing led to a decrease in VIRMA expression, while its overexpression resulted in its upregulation (Figure 5). Other complex proteins remained unaffected despite Hakai’s modulation. These results partially contrast with previous studies in *Drosophila* and human HeLa and U2OS cells, where Hakai was shown to form a stable complex with WTAP, ZC3H13 and VIRMA and its depletion led to a marked reduction not only in VIRMA protein levels but also in WTAP and ZC3H13. In those models, Hakai was proposed to act as a stabilizing factor for these “orphan” proteins (subunits that become unstable and are targeted for degradation in the absence of their binding partners) [31,35]. This context-specific dependency may be explained by the recently proposed flexible architecture of the MACOM complex [11]. Only three components (WTAP, VIRMA, and ZC3H13) of the complex were found, while Hakai, despite being present in the SDS-PAGE analysis, was missing in cryo-EM structure. In this context, Hakai could be adopting a flexible conformation, being dispensable for RNA substrate binding and for methyltransferase activity [11]. Given that Hakai functions as an E3 ubiquitin-ligase, we explored whether its ubiquitin-ligase activity might contribute to the regulation of the m^6^A writer complex. Using Hakin-1 (specific inhibitor of Hakai ubiquitin-ligase activity) [19], we found that the catalytic inhibition of Hakai did not alter the protein levels of VIRMA or any other components of the m^6^A complex analysed. Moreover, a CHX chase assay under control conditions and upon Hakai overexpression excluded a VIRMA-stabilizing role for Hakai (Figure 5). This suggests that Hakai does not exert its function within the m^6^A writer complex through its canonical E3 ubiquitin-ligase activity and that instead, its role may be mediated through non-catalytic mechanisms. This is consistent with previous reports in *Drosophila*, where Hakai stabilizes three components of the complex with no requirement of its ligase activity [35].

Taking into account that m^6^A methylation takes place in the nucleus and that silencing of Hakai led to reduced m^6^A levels without affecting METTL3 protein expression, we hypothesized that Hakai may influence METTL3 subcellular localization, a reported mechanism for other members of the complex. In particular, the mouse protein Zc3h13 (homolog to human ZC3H13) is required for the nuclear retention of mouse proteins Wtap, Virilizer and Hakai, as well as Metll3 and Metll14, without altering total protein levels [36]. Our data shows that Hakai favours the nuclear retention of METTL3, with its absence leading to altered compartmentalization, which may underlie the decreased global m^6^A methylation levels. We also explored if Hakai silencing would result in a widespread decrease in m^6^A methylation across target transcripts. However, our MeRIP-seq analysis in the HT29 cell line not only revealed hypomethylation of 31 mRNAs, but also hypermethylation of 25 mRNAs (Figure 2). These results are consistent with previous observations involving m^6^A writer components such as ZC3H13, WTAP, or VIRMA, where silencing predominantly resulted in hypomethylated mRNAs, yet hypermethylation at specific mRNAs was also observed. The opposite phenomenon has been reported for m^6^A erasers like FTO, where gene deletion resulted in a expected global hypermethylation, but also causing hypomethylation in a subset of mRNAs [36–39].

Functionally, methylated mRNAs identified by MeRIP-seq were predominantly associated with pathways involved in the immune cell recruitment, cytokine signalling and inflammation. While no previous reports directly link Hakai to the specific genes identified in our dataset, several of these genes have been identified as m^6^A methylation targets in various physiological and pathological contexts [40,41]. Moreover, correlations between the CBLL1 gene (Hakai protein) and immune cell infiltration in several diseases have been identified, including various types of cancer such as hepatocellular carcinoma and lung cancer [42–46]. Supporting this findings, Hakai is reported to influence the immune microenvironment in periodontitis through regulation of TNF family members and cytokines [47]. However, the immunoregulatory effect of Hakai appears to be context-dependent. For example, in endometriosis, a negative correlation between CBLL1 and M2 macrophages was reported, suggesting an immunoprotective role; whereas in ischemic stroke a positive correlation was observed, implying a pro-inflammatory role [48,49]. These findings and our own results suggest that Hakai may participate in the regulation of the crosstalk between tumour cells and the immune microenvironment, contributing either to immune suppression or immune activation depending on the cellular context and pathophysiological conditions.

We further sought to characterize the effects of m^6^A methylation on Hakai target genes identified by MeRIP-seq. Hakai silencing led to an increased expression of key immune-related genes, whereas Hakai overexpression resulted in their downregulation (Figure 3). These genes also showed increased m^6^A methylation upon Hakai silencing, suggesting a positive correlation between m^6^A levels and transcript levels. Treatment with the global methylation inhibitor 3-DZA partially reversed the observed effects of Hakai silencing, supporting that the process may be at least partially driven by m^6^A-dependent methylation mechanism. Our experiments assessing mRNA stability with ActD suggests that the expression changes are unlikely the results from altered transcript stability. Instead, our data are more consistent with posttranscriptional mechanisms accounting for the observed differences. Collectively, the data suggests that through m^6^A modulation, Hakai influences the expression of immune-regulatory genes that could affect immune cell recruitment and activation, shaping the tumour microenvironment in CRC. In this context, m^6^A reader proteins such as YTHDF1 have been proven particularly important, promoting the translation of methylated transcripts without altering their stability [50,51].

Based on our findings showing that Hakai regulates several immune-related genes in CRC cells and considering recent studies linking m^6^A methylation to modulation of the tumour microenvironment, we explored Hakai’s role in immune cells. We examined whether the gene expression changes induced by Hakai modulation in CRC cells were also observed in PBMCs. Interestingly, Hakai overexpression increased IFNB1 expression only in PBMCs, whereas it reduced IFNB1 levels in HT29 cells (Figure 5). This reinforce the notion that Hakai’s regulatory role is context-dependent and varies between cell types [51,52] Given these differences between tumour and immune cells, we next explored whether factors secreted by Hakai-silenced tumour cells could affect peripheral and gut immune cell responses. Using conditioned media from tumour cells to treat PBMCs and LPMCs, we observed increased expression of inflammatory and cytotoxic markers, notably IFNG and cytotoxic effectors, which are associated with enhanced antitumour immunity and favourable prognosis in several cancers, including CRC [52–57]. Thus, Hakai appears to favour an environment that suppresses cytotoxic immune activity, with its silencing leading to a more immunostimulatory and potentially antitumoural profile. Given that epitranscriptomic alterations can reshape the tumour secretome, influencing immune cell function [58–60], our results suggest that Hakai not only modulates m^6^A-dependent gene expression in tumour cells, but also indirectly affects immune landscape through secreted signals. While the precise mechanisms remain to be fully elucidated, our data poses Hakai as a potential central node in the crosstalk between tumour cells and the immune microenvironment. These findings position Hakai as a potential mediator of tumour–immune crosstalk and a target for future investigation in antitumour immunity.

## CONCLUSIONS

In this work, we investigated Hakai’s involvement in transcriptomic and epitranscriptomic regulation, which has been largely overlooked compared to its well-known function as an E3 ubiquitin ligase regulating protein stability. Its overall impact on the transcriptome of colorectal cancer cells, both in monolayer and in 3D tumourspheres is limited. However, we found a significant impact on the overall m^6^A methylation levels of colorectal cancer cells. Mechanistically, we found an interaction with the MACOM protein VIRMA. Hakai regulated VIRMA protein levels independently of its E3 ubiquitin-ligase activity or post-translational stabilization. Moreover, Hakai silencing also altered the localization of METTL3, the catalytic subunit of the MAC complex. Methylome analysis revealed changes in genes related to immune responses, suggesting a previously undescribed role for Hakai in this process. To explore this further, we silenced Hakai in colorectal cancer cells and exposed patient-derived immune cells to their conditioned medium. This led to modulation of several immune response markers in these immune cells. Overall, our findings position Hakai at the intersection of m⁶A methylation control and tumour-immune response, and suggest that Hakai silencing could enhance antitumour immunity in colorectal cancer. However, further studies are needed to clarify the complex, context-dependent mechanisms linking m⁶A regulation and immune responses in the tumour microenvironment.

## DECLARATIONS

### Ethics approval and consent to participate

Human colon samples were collected by the Department of Pathological Anatomy of the Tor Vergata University Hospital (Italy), with informed consent from the patients. Studies involving human samples in Italy were reviewed and approved by the Ethical Review Committee of the Tor Vergata University Hospital (experimentation register no. 42/12). Additionally, human primary tumour samples were collected from General and Digestive Surgery Service of the A Coruña University Hospital (Spain), with informed consent from the patients. PBMC from healthy donors were also collected at the Galician Blood, Organs and Tissues Agency (ADOS, Spain). Studies involving human samples in Spain were reviewed and approved by the Research Ethics Committee of A Coruña-Ferrol, in accordance with the ethical procedures established and described in the “Organic Law on Biomedical Research” of July 14, 2007, under Spanish regulations (code of ethical protocols: 2021/130 and 2024/217).

### Consent for publication

Not aplicable

### Availability of data and materials

The datasets used and/or analysed during the current study are available from the corresponding author on reasonable request.

### Authors’ contributions

Experimental procedures were performed by MQ and AI. Bioinformatic analyses were conducted by VS and MQ. Writing of the original draft was carried out by MQ and AF. Contributions to writing, revision, and editing of the manuscript were made by AF, MQ, JJER, and IM. Experimental design and project conceptualization were conducted by AF and IM. Project conceptualization and funding acquisition were led by AF. All authors read and approved the final manuscript.

## Acknowledgements

The authors would like to acknowledge Dr. Yasuyuki Fujita (Hokkaido University, Japan) and Dr. Jianzhao Liu (Life Sciences Institute, Zhejiang University, China) for kindly providing plasmids used in this work.

## LIST OF ABBREVIATIONS

3-DZA: 3-(deazaadenosine)
ActD: Actinomycin D
CHX: Cycloheximide
CRC: Colorectal cancer
DDX39B: DEAD box helicase 39B
DEGs: Differentially expressed genes
FDR: False Discovery Rate
HYB: Hakai-pY-binding domain
IgG: Immunoglobulin G
LFC: Log-fold scale
LPMCs: Lamina propria mononuclear cells
m^6^AN6: methyladenine
MAC m^6^A: METTL Complex
MACOM: m^6^A METTL-associated complex
MeRIP: Seq Methylated RNA Immunoprecipitation Sequencing
METTL3: Methyltransferase-like 3
METTL14: Methyltransferase-like 14
NK: Natural Killer
PBMCs: Peripheral blood mononuclear cells
PCA: Principal component analysis
PPI: Protein-protein interaction
RNA-seq: Ribonucleic acid sequencing
RT-PCR: Reverse transcription polymerase chain reaction
SAM: S-adenosylmethionine-dependent methyltransferases
TME: Tumor microenvironment

